# Poor man's two photon imaging: Scanning Laser Upconversion Microscopy

**DOI:** 10.1101/138909

**Authors:** Cecilia Sorbello, Roberto Etchenique

## Abstract

Two photon microscopy is one of the most powerful techniques for optical images. Its deep penetration, low scattering and mainly its sectioning power allows exquisite control on all three axes of 3D samples, both for imaging and actuation. Unfortunately, this technique involves very expensive femtosecond lasers. Upconverting Nanoparticles display multiphoton absorption at low instantaneous power, which can be achieved by means of cheap laser diodes. Scanning Laser Upconversion Microscopy offers the deep penetration and low scattering of NIR excitation together with the sectioning power of multiphoton techniques for under US$ 1000.

## Introduction

Multiphoton microscopies offer important advantages over their 1-photon counterparts: NIR excitation, deeper penetration in living tissue, lower scattering and z-axis sectioning. Among them, the direct sectioning power that arises from a nonlinear emission-excitation relationship is perhaps the more interesting one. The fact that the emission follows the square (or a superior power) of the excitation makes it high just in the focal zone, leaving unperturbed the regions below and over the focal plane, especially when high aperture objectives are used. Optical sectioning can also be obtained through confocal microscopy. However, while confocal techniques illuminates a full bicone of light and pick the chosen plane by means of the collection optics (pinhole), multiphoton microscopies produces the very excitation just in the focal plane, preventing the rest of the sample from reaching excited states that led to emission or further photochemical pathways. This characteristic has two additional advantages: a) photobleaching of probes is confined to the focal zone and b) photochemical actuators can be activated having real 3D positioning.

Since Denk reported the first two-photon microscope^1^, many probes and actuators have been designed and developed for imaging^2–4^ and triggering biological systems^5–8^. Two (or more) photon absorption is a hard way toward excitation. Most substances have negligible absorption for processes that involve more than one photon. In order to obtain a reasonable emission flux after two-photon (2P) excitation, an enormous instantaneous power density must be delivered into the sample at the time that the average power is kept low in order to prevent damage through overheating. Ti-Sapphire lasers are delicate, big and expensive, being these the main reasons because multiphoton microscopies are not so widely extended as their advantages suggest. In a typical application a 4W average power Ti-Sapphire laser, 140 fs long pulses, 80 MHz repetition can conduct a maximum of 4.5×10^13^ W/cm^2^ over the sample^9^.

Luminescent nanoparticles are established as useful tools for imaging biological systems. Their uses range from robust probes with very high tolerance to photobleaching^10^ to nanosized barcoding systems^11^. Among nanoparticles, lanthanide-containing nanocrystals show a rather odd behaviour: upconversion. Upconversion is a photophysical process by which two or more photons of a lower energy are absorbed by a system that eventually decays through the emission of a higher energy photon. Multiphoton excitation and upconversion are two quite different processes. In the first mechanism, two or more photons are absorbed quasi-simultaneously, populating the emitting excited state from the ground level. A formalism having a virtual state is often used to depict this process, as shown in Figure 1a. Conversely, upconversion involves long lived real states that lays between the ground and the emitting states, which can be populated, accumulating energy that eventually will be released as emission of a short wavelength photon. Several internal mechanisms are usually present in upconverting systems, Fig 1b and 1c depict the more widely studied mechanisms: Excited State Absorption (ESA) and Energy Transfer Upconversion (ETU).

**Figure 1.**
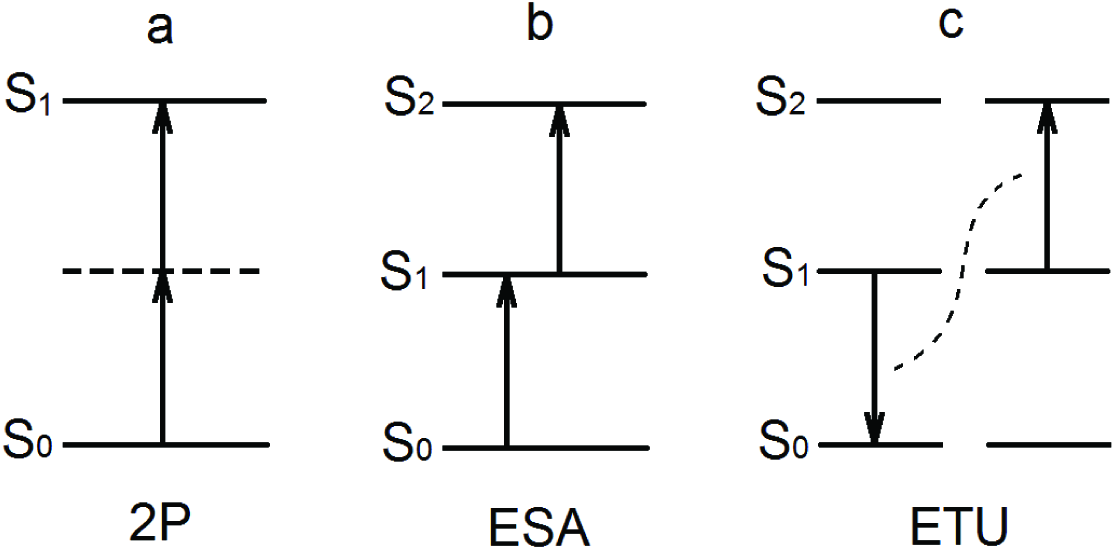
Scheme of the mechanisms of multiphoton absorption and upconversion. a) Two photon absorption, depicting the virtual state between S1 and S2. b) Excited state absorption, showing the real intermediate state. c) Energy transfer upconversion.

The presence of this intermediate energy states has a profound effect in the absorption cross section. While usual 2P absorption needs both photons to be absorbed within the time of a transition (~10^−15^s) and therefore very high instantaneous powers, upconversion mechanisms operate in the range of 10^−5^ s and thus they can be efficient even at excitation powers of ten orders of magnitude lower than 2P. High efficiency upconverting phosphors can present visible emission under excitation as low as fractions of W/cm^2^. In other words, while a good 2P emitter can have a cross section of hundreds of GM (1 Goeppert Mayer = 10^−50^ cm^4^ s), a typical UCNP can exhibit values higher than 10^8^ GM^12^.

Even though the photophysics underlying upconversion is different from that of 2P absorption, the equations governing both mechanisms are essentially the same. In stationary state, the emission intensity will scale as a power of the excitation power, where the power is equal to the number of photons needed to reach the emissive excited state. Due to this fact, it is expected that upconversion microscopy would behave as usual multiphoton microscopies, yielding z-axis sectioning. However, reports on using upconverting probes to take advantage on their intrinsic sectioning capabilities are scarce and negative. Yang et.al. ^13^ have used upconversion nanoparticles (UCNPs) in a scanning scheme to image labelled cells, although they have not succeeded in obtain intrinsic sectioning power, attributing this failure to the nature of upconversion mechanism and using a confocal pinhole to achieve z-sectioning. Veggel^14^ have pointed out the main problem of upconversion excitation: The same characteristics that allow multiphoton absorption with high cross section at low power densities imply that the saturation of the intermediate states becomes important at rather low excitation fluxes, linearizing the effective emission-to-excitation dependence. Some authors have tried to circumvent this issue by applying confocal techniques to upconversion. This is a hard task to accomplish due to the long characteristic times of UCNP emission, which implies very low scanning sweeping speed or image smearing. Romanowsky have devised a clever procedure to reconstruct smeared images through deconvolution, useful for sparse labelling^15^, while Pierce have attacked the problem by means of line confocal microscopy^16^, where just the slow axis needs to be swept. However, these latter approaches also allow image sectioning by means of confocal procedures, not using the intrinsic sectioning power of upconversion as a nonlinear emission process.

We present in this work the first demonstration of full multiphoton sectioning obtained by using upconversion nanoprobes. We characterize the method, showing that the only requisite for having good sectioning power is to prevent saturation, being upconversion similar to traditional multiphoton microscopies in this respect. We measured the main optical characteristics of upconversion microscopy at different regimes. Finally, we solved the problem of having simultaneously long collection times and fast scanning by using parallel descanning through a CCD sensor, and obtained optical sections of highly homogeneously labeled specimens using a low power laser diode as source and an overall equipment three orders of magnitude cheaper than the usual 2P microscopes. In this way, Scanning Laser Upconversion Microscopy (SLUM) opens a new path to inexpensive but precise imaging and/or manipulation of living objects.

## Results and discussion

Er-containing UCNPs ((NaYF_4_: 2mol% Er^3+^: 30mol% Yb^3+^) present absorption at NIR (980 nm) and anti-stokes emission at two main bands: 500-550 nm ((^4^S_3/2_-^4^I_15/2_ and ^2^H_11/2_-^4^I_15/2_ transitions) and 640-670 nm (^4^F_9/2_-^4^I_15/2_). The emission spectra is given as supplementary information (Figure S1). Both green and red emission bands correspond to biphotonic processes. Therefore, in absence of saturation the emission scales as the square of the excitation intensity. Upconverting emission is a rather slow process, with characteristic times in the hundreds of µs, which are dependent on the exact composition of the UCNPs. Figure S2 shows the temporal characteristics of the UCNPs used in this work. The long characteristic times associated imply that any imaging based on scanning must be conveyed in one of the following ways: 1) performing slow scan procedures, to collect all the light from any point before passing to the next, or 2) using a parallel descanning method capable to get the emission from points that are not longer excited, without smearing the image by attributing this light to the point being scanned at the moment of detection. The first procedure would make the scan process very low. At 1 ms/pixel, in order to collect most upconverted photons, a VGA-sized image would take 5 minutes. Parallel descanning can solve this issue. We have chosen to use a CCD camera as parallel descanning method. In this way, any pixel at the CCD sensor will accumulate all the photons during a line or frame scan, while the scanning beam can be moved very fast through the specimen. The diagram of the setup is given as supporting information (Figure S3). This fast scanning is also key to prevent saturation of the probes and therefore keep z-axis sectioning while maintaining high average excitation power (*vide infra*).

In order to evaluate the sectioning capabilities of SLUM it is convenient to image a simple object. We chose a thin homogeneous layer of UCNPs embedded in polystyrene by spin coating. For a Gaussian beam focused onto a sample, the lateral distribution of intensity at the focal plane can be describes as:

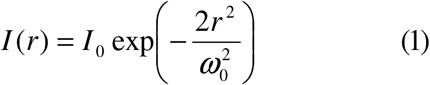

where I_0_ is the intensity at the center of the distribution, r is the distance from the propagation axis and ω_0_, the Gaussian beam waist, is the distance at which the intensity has decreased to 1/e^2^≅ 0.135.

The waist ω_0_ is related with the wavelength λ and the angular aperture θ:

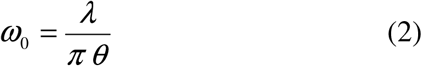

On the propagation axis, the size of the waist increases below and above the focal plane:

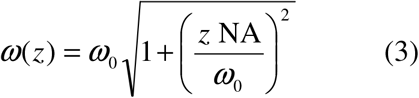

where NA = n sin *θ* is the numerical aperture of the objective (n = refractive index of the sample). Therefore, the equivalent area of the illuminated zone at any distance *z* from the focus is:

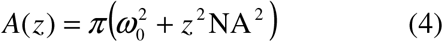

For a given photon flux and in absence of absorption, equation (4) represents the reciprocal of the excitation density. For a laterally infinite and homogeneous emissive thin layer which is scanned in x and y directions, the total collection light does not depend on the focusing, but only on the emission intensity of the layer. Therefore, if this emission is due to a linear process, no change will be apparent by changing z-position. However, for a nonlinear process, the collected emission of the layer will be a function of the position as follows:

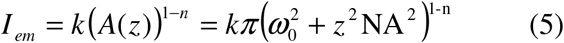

where *k* is a constant that includes all instrumental parameters and also the emission quantum yield of the probe, and n is the number of photons of the nonlinear process. For a typical 2P process, n=2, while usual upconversion mechanisms often provide fractional numbers, indicating a multiplicity of pathways by which the emissive state can be reached.

Figure 2a shows the experimental results for the imaging of a thin sheet of UCNPs embedded into a polystyrene film at two different excitation intensities. The emission increases at the focal plane as expected for a multiphoton process.

**Figure 2:**
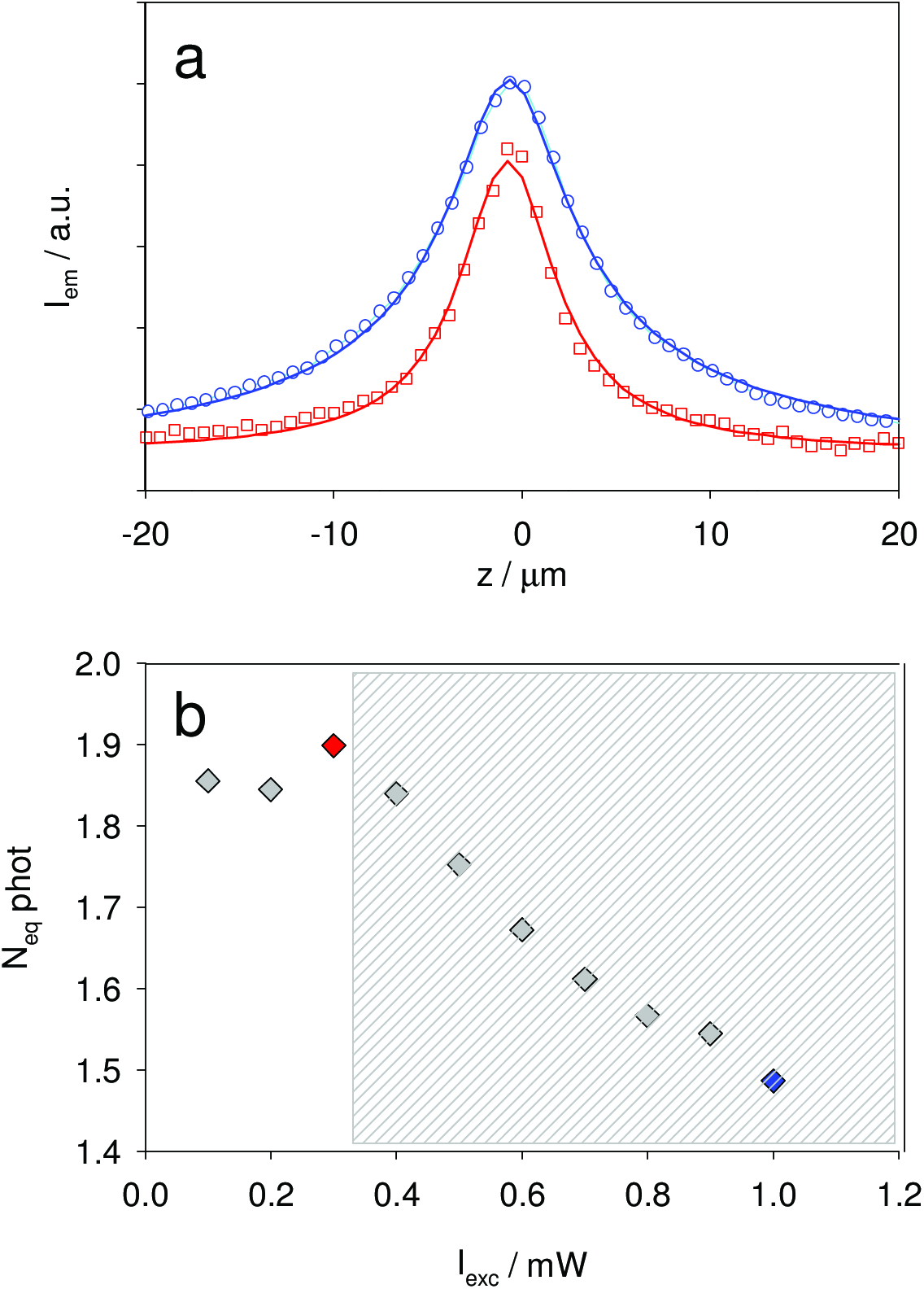
a) Performance of SLUM z-sectioning at two different excitation intensities. The specimen is an infinite thin sheet of upconverting nanoparticles. P = 0.3 mW (red circles) and P = 1 mW (blue squares), both at a scanning speed of 9.6 cm/s. The solid lines are the best fits to the equation (5) b) Dependence of the equivalent number of photons with the excitation power. The red and blue points correspond to the curves in (a). The zone where focal saturation is present is grayed.

For low excitation intensity (100-400 μW), at a scanning speed of 9.6 cm/s, the number of photons that best fit the experimental data is near 1.90, indicating that a 2-photon process is the main emission pathway. In this conditions, the z-range in which the emission of the layer diminishes to a half is about 6.6 μm at half height (Figure 2a, squares). At a higher intensity (1 mW, circles), the curve broadens as expected from a higher contribution of 1-photon excitation due to saturation near the focal zone. The relationship between excitation intensity and photonicity, showing the saturation zone, is depicted in Figure 2b.

These results clearly show that saturation must be avoided to get good sectioning power. There are two ways to achieve this goal: to lower the excitation density or to increase the scanning speed, in order to allow the probes to receive light just a very small fraction of the time. As was already demonstrated, both strategies are completely equivalent and short pulse excitation yield the same results that low intensity in order to prevent saturation^12^.

The Point Spread Function (PSF) of an imaging system describes the response of this system to a point object. Any obtained image can be described as the convolution of the imaged object with the PSF of the system. Therefore, it is useful to obtain the PSF of a system as a basis for comparison^17^. To estimate the PSF of the SLUM at different saturation conditions , we have focused an stationary beam on a thin layer of UCNPs, forming an emitting object with negligible size in z-direction and near-Gaussian profile in x and y directions. The obtained raw images were deconvolved with the estimated size of the emitting object and the results at different excitation intensities are depicted in figure 3.

**Figure 3:**
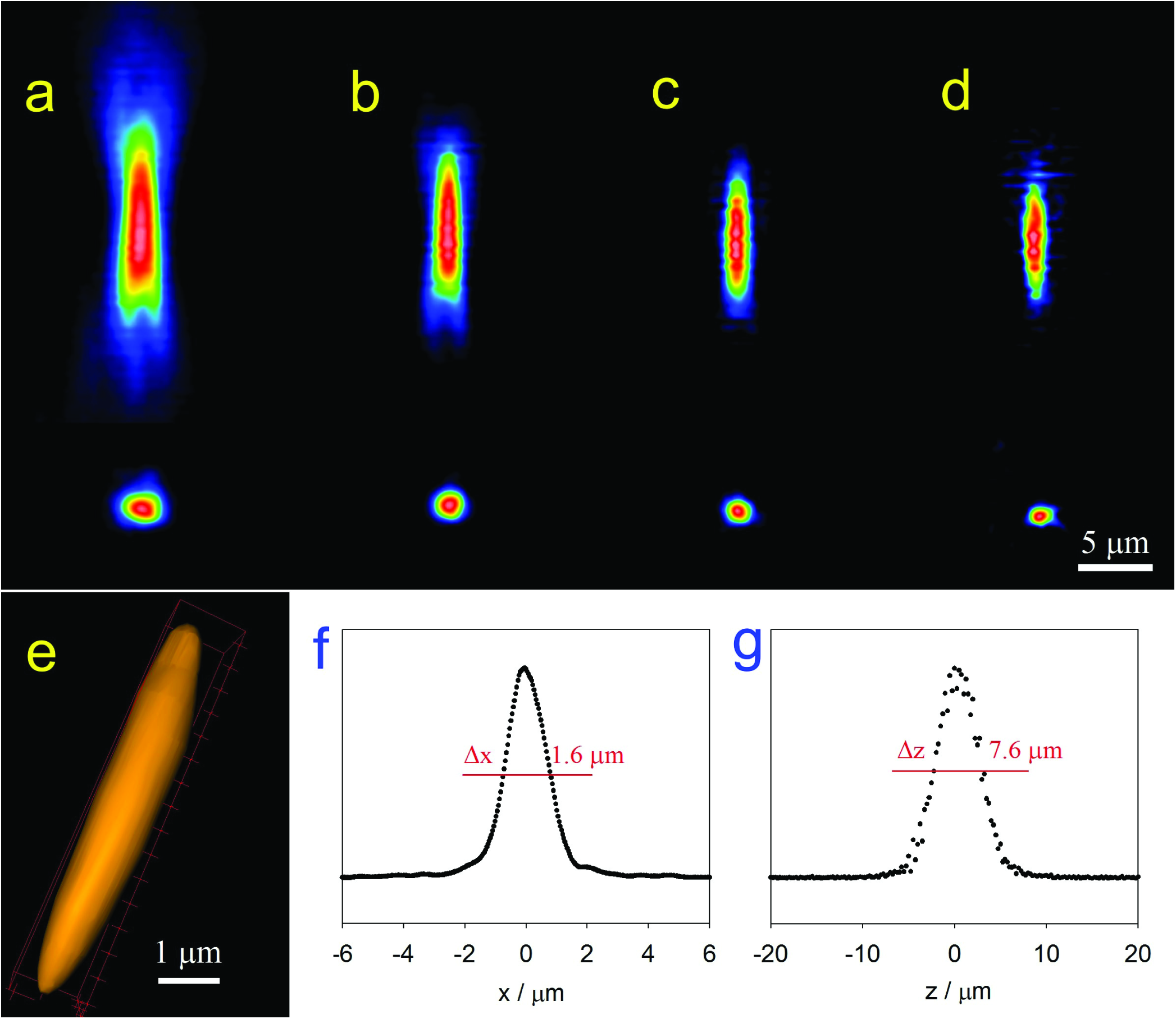
a) False colour z-x planes of the point spread function of SLUM at different excitation intensities. a) 3 mW, b) 100 µW, c) 25 µW, d) 10 µW (NA=0.65). e) 3D reconstruction of the PSF at panel c (p = 25 µW). f) Resolution of SLUM in lateral (x,y) directions. g) resolution of SLUM in axial (z) direction (p = 25 µW).

While the excitation power does not substantially change the on-plane resolution (x,y), it has a profound effect in the axial (z) direction. At high excitation density, saturation of intermediate states allow the emitters to bright through a 1-photon linear regime. At this conditions the sectioning power of the SLUM vanishes, showing the bi-conical excitation as a normal epifluorescence microscope. On the contrary, at lower intensities, nonlinear multiphoton regime takes place and the PSF shows the typical z-localized ovoid shape of 2P microscopies. For excitation densities below 25 mW (~1200 W/cm2) and using an objective of NA = 0.65 the lateral resolution is Δx = 1.6 µm and the axial resolution is Δz = 7.6 µm. The differences with the theoretical values for perfectly focused Gaussian beam (Δx =0.9 µm and Δz = 6.9 µm respectively) can be ascribed to chromatic aberration and slight misaligning through the optical system.

In order to show the power of the technique, we have chosen a suitable specimen to compare z-sectioning. Figure 4 shows series of images of a pollen grain (*Abutilon grandifolium*) covered with UCNPs and immersed into ethyl salicylate (η = 1.52) which was imaged at different planes with the same optics but varying the parameters in order to obtain linear or multiphoton images. Seven of these image planes are depicted in the panel (a). The row (b) shows the specimen as seen through bright field (λ = 525nm) at the chosen focused planes. As usual, out-of-focus light obscure the fine details of the specimen. Linear emission microscopy is not capable to solve this issue: The row (c) shows the image under Koehler epi-illumination, where NIR (980 nm) excitation intensity independent on the z position. Although the upconverted emission is a multiphoton process, as the excitation density does not change at different planes, the overall effect is the same as if where imaged by means of conventional 1-photon fluorescence, as usual in upconversion microscopies. The collected emission of every plane throughout the sample have a similar bright, which is given by the excitation intensity. Therefore, the out of focus light also prevent any fine structure of the grain to be detected. On the other hand, the sectioning power of SLUM is clearly depicted in Figure 4d: The images were taken with p=0.5 mW excitation power at a scanning speed of 19.2 cm/s. At this speed, and considering the PSF, the focal volume is excited during about 8.3 µs during the beam flight. The equatorial image shows that the non labelled interior of the grain is clearly observed, even through the bright exine capsule. A reconstructed 3D view of the pollen grain is also shown (Figure 4e), in complete accordance with the image obtained by SEM (Figure 4f). While the SEM picture is somewhat distorted due to the high vacuum needed to image it, the 3D reconstruction preserves the near spherical shape.

**Figure 4:**
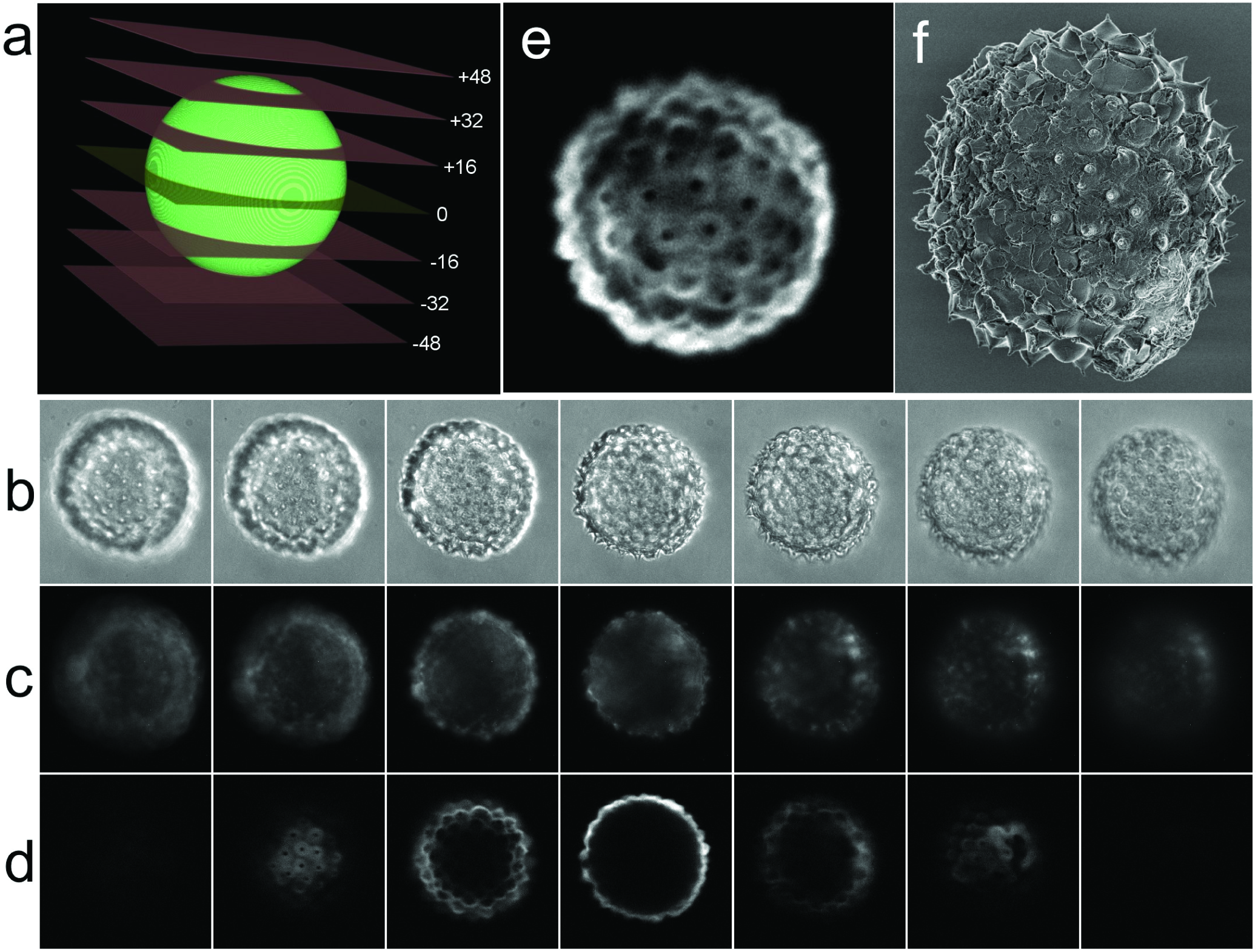
a) Scheme of the depicted focal planes in panels b-d. Distances from the equatorial plane are given in µm. b) Bright field images of a pollen grain (*Abutilon grandifolium*) illuminated with a 525 nm LED source. c) Linear epi-illumination images of the same specimen using a non focused 980nm Laser Diode excitation. d) SLUM images at p= 500 µW, v = 19.2 cm/s of the same specimen showing 2-P sectioning power. All images were taken through a 40×0.65 objective. e) 3D reconstruction of the pollen grain. f) SEM image of the pollen grain in vacuum.

It is important to emphasize that the observed sectioning was not obtaining by means of eliminating off-focus light as in confocal procedures, but due to the intrinsic nonlinear nature of the involved process. This fact indicates that the same strategy can be used to achieve true 3D localized excitation than can be used to elicit photochemical responses (i.e. drug uncaging, polymerization, etc.) in thick systems, as usual with standard 2P techniques. Further research on these applications are being carried on.

## Conclusions

We have shown that true z-sectioning can be performed using upconverting nanoparticles as probes. The key of this achievement lays on the use of very low duty cycle pulse excitation that prevents saturation of the intermediate states of the UCNPs, thus keeping the emission in nonlinear multiphoton regime. This method presents figures of merit near to the theoretical ones for the used optics and allows to build a very low cost 2P microscope, where solid-state laser diodes replaces the bulky and expensive Ti-Sapphire femtosecond oscillators. We have tested a simple version of such microscope as a proof of concept to show sectioning in a spheroidal pollen grain. Given that the multiphoton sectioning is intrinsic to the nonlinear excitation mechanism, the same technique could be used to perform photolithography and phototriggering, opening a wide field of low cost and small-sized methods for sensing and actuating with exquisite 3D resolution.

## Acknowledgements

This research was supported by the National Agency for Science and Technology Promotion, CONICET, and the University of Buenos Aires R. E. is a member of CONICET.

---

**Figure S1:**
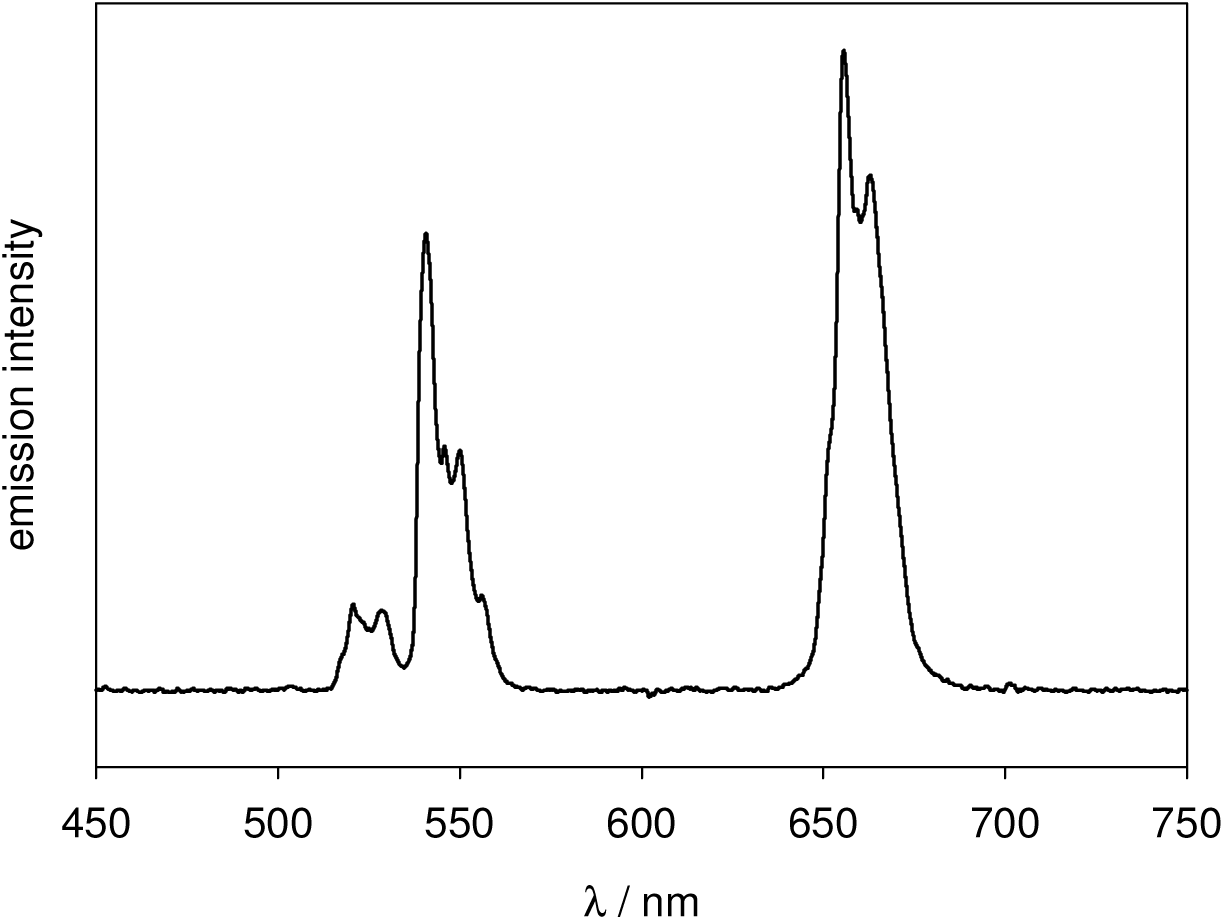
Emission of the used UCNPs under irradiation at 980 nm.

**Figure S2:**
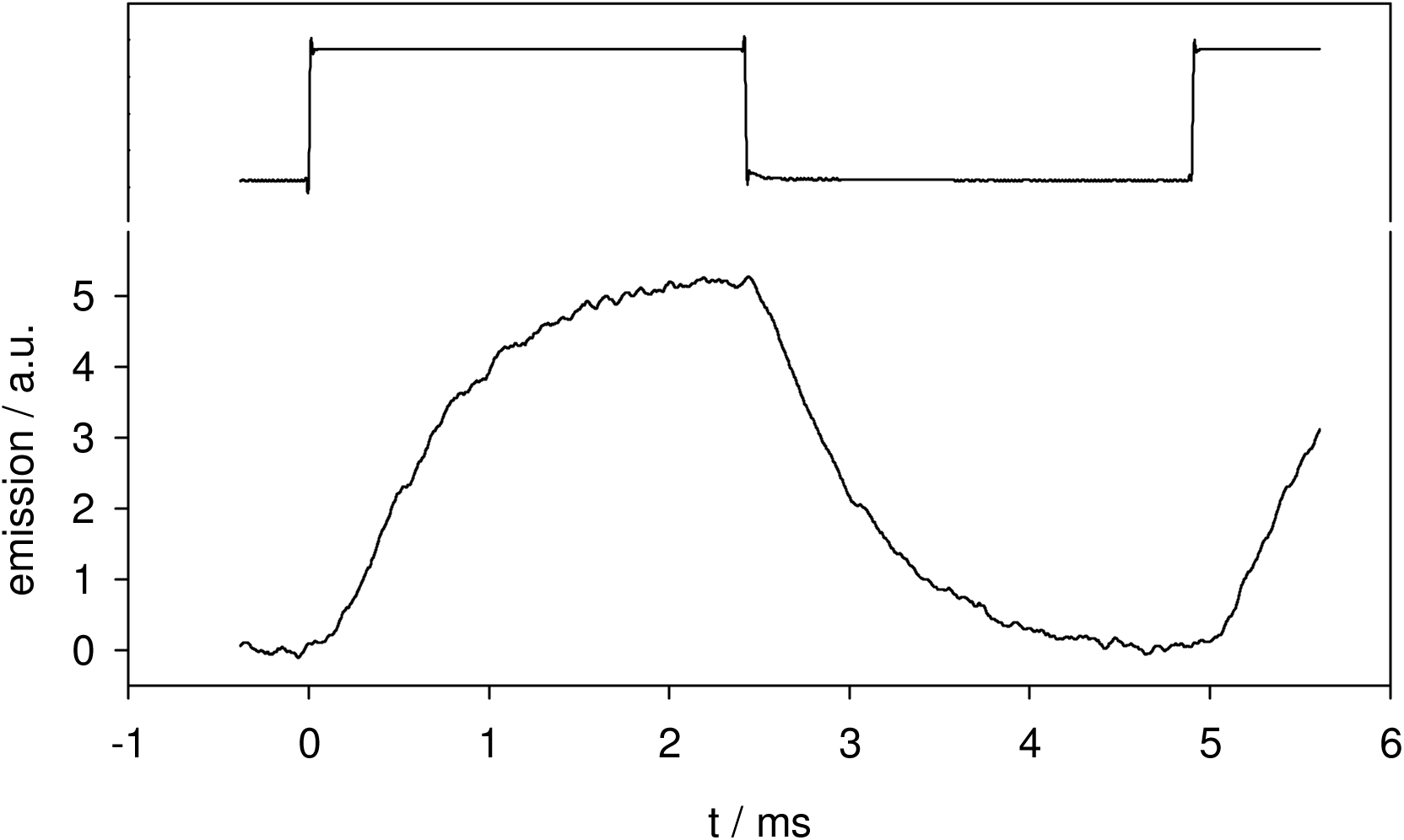
Top: Irradiation of the 980 nm laser (ON-OFF). Bottom: Time-resolved emission of the used UCNPs.

**Figure S3:**
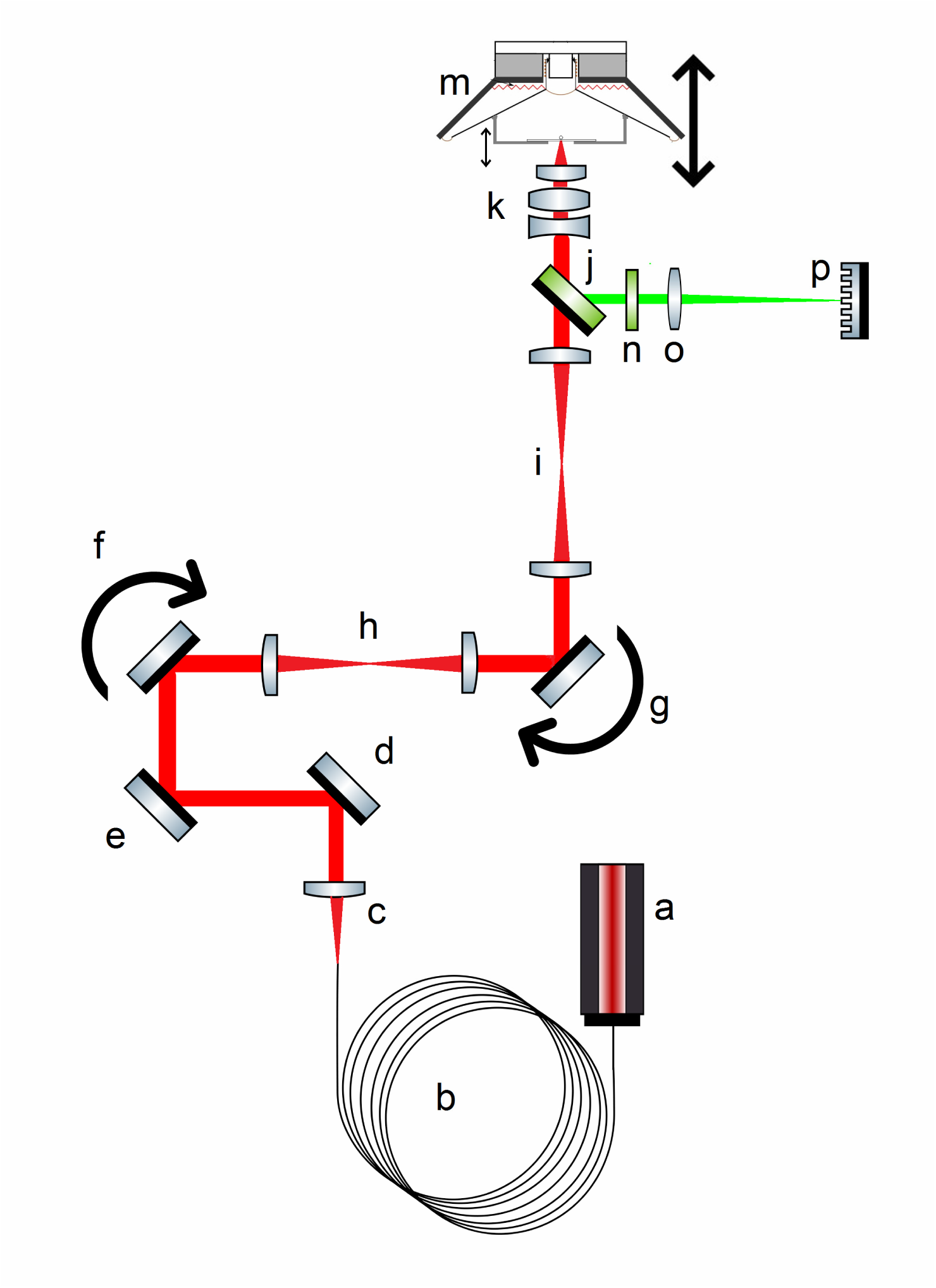
Overall scheme of the SLUM. A NIR (980 nm) solid state 10 mW laser (a) is coupled to a single mode fiber optics (b), collimated with plano convex lens and sent using two mirrors (d,e) to a pair of X-Y scan galvos (f,g) which are confocal through a telescope h. The scanned NIR beam is enlarged with a second telescope (i), pass through a dichroic mirror (j) and directed to the objective (k) to the specimen which z-position can be modified with a speaker (m). The emission of the specimen is collected from the objective and reflected by the dichroic mirror, pass through a block filter that removes any residual NIR excitation (n) and focused through a lens tube (o) to the CCD sensor (p).

